# Focal versus distributed temporal cortex activity for speech sound category assignment

**DOI:** 10.1101/133272

**Authors:** Sophie Bouton, Valérian Chambon, Rémi Tyrand, Adrian G. Guggisberg, Margitta Seeck, Sami Karkar, Dimitri van de Ville, Anne-Lise Giraud

## Abstract

Percepts and words can be decoded from largely distributed neural activity measures. The existence of widespread representations might, however, conflict with the fundamental notions of hierarchical processing and efficient coding. Using fMRI and MEG during syllable identification, we first show that sensory and decisional activity co-localize to a restricted part of the posterior superior temporal cortex. Next, using intracortical recordings we demonstrate that early and focal neural activity in this region distinguishes correct from incorrect decisions and can be machine-decoded to classify syllables. Crucially, significant machine-decoding was possible from neuronal activity sampled across widespread regions, despite weak or absent sensory or decision-related responses. These findings show that a complex behavior like speech sound categorization relies on an efficient readout of focal neural activity, while distributed activity, although decodable by machine-learning, reflects collateral processes of sensory perception and decision.

## Introduction

Does all the information that is encoded in our brain contribute to our representations and our decisions? The discovery of spatially distributed cortical representations, exploitable for “mind reading” in all domains of cognitive neuroscience during the past decade (1–5) raises fundamental issues about the nature of neural coding in the human brain. These findings even led scientists to reconsider the notion of local computational units, such as canonical microcircuits (6,7). Broadly distributed representations, as recently exemplified with the extreme scattering of word meaning representations throughout the brain (8), must, however, be interpreted with caution. Such data-driven results could lead us to conclude that these representations are useful when performing cognitive operations, when in fact they might only follow from these operations and reflect associative processes. Spatially-distributed patterns could be taken to indicate that the information they contain is critical to cognitive operations, as for instance in simple stimulus categorization (9). Yet, there is no principled *a-priori* reason why the sensitivity of techniques probing multidimensional neurophysiological data, e.g., machine decoding, would reflect the capacity of our brain to use broadly distributed neural patterns for specific cognitive operations (10–12).

Interpreting broadly distributed spatial maps for speech sounds can be particularly difficult. Unlike visual stimuli, whose identity relies heavily on spatial encoding, speech sound identity is embodied in a temporal form, and mainly relies on temporal encoding (13,14). Despite the importance of hierarchical temporal processing in speech, large cortex coverage with fMRI and more recently with ECoG was used to demonstrate 1) that the original acoustic speech signal can be reliably reconstructed from broadly distributed high frequency neural activity sampled cross-regionally throughout the superior temporal lobe (15–17), and 2) that local phonemic identity information in speech is poorly encoded by temporally resolved neural activity (2) but finely represented by distributed cortical patterns covering a significant portion of the left temporal lobe (1). Because optimal decoding occurs when redundant information from contiguous but functionally distinct territories is pooled together, assigning perceptual relevance to such large-scale representations ultimately conflicts with the notion that speech sounds are first spectro-temporally encoded in auditory cortex, before being more abstractly recoded in downstream areas (18). Accordingly, focal lesions of the temporal lobe can selectively impair different speech perception processes (19), and recent studies in monkey even show that auditory decision-making causally relies on focal auditory cortex activity (20).

In summary, although multivariate pattern analysis and related decoding techniques have become popular (and prevalent) in the systems neuroscience literature, they may lead to misleading interpretations. This is because demonstrating that a particular stimulus attribute or category can be decoded from regional neuronal activity does not mean that the region is performing any decoding or categorical processing. In other words, there is an important conceptual leap between machine and brain decoding of neuronal activity. As an extreme example, phonemes could be classified using a machine-learning scheme applied to primary sensory afferents from the ear. However, this does not mean that the brain has yet decoded these signals – just that there is, by definition, sufficient information in this auditory input to support subsequent hierarchical decoding. To address this issue, we distinguished between the ability of a classifier to decode the stimulus category from neuronal responses at various levels in the auditory hierarchy and the ability of a linear model to estimate from neural responses the perceptual processes in a speech sound category assignment paradigm. In brief, we found that speech sound category assignment relied on neural activity present in a circumscribed part of the auditory hierarchy, at particular peristimulus times. In contrast, multivariate machine decoding returned positive results from a large brain network including regions where no evoked activity could be detected.

## Results

We first explored explicit phoneme recognition using a simple syllable categorization task and measured global neural activity with fMRI and MEG in respectively 16 and 31 healthy volunteers (see methods and S1 Text). The subjects had to decide which syllable they heard in a /ba/ /da/ continuum, where the onset value of the second formant (F2) and the F2-slope linearly covaried in six steps (Fig 1A). These two first experiments served to delineate at the whole brain level those brain regions that were sensitive to 1-linear variations of F2 and 2-perceptual decisional effort as assessed using behavior-based negative *d’* values (Fig 1B, 2A) (see methods and S1 Text). Critically, because the slope of the 2^nd^ formant is steeper for the /da/ than for the /ba/ phoneme, we expected the /da/ stimulus to activate a larger cortical surface than the /ba/ stimulus, and hence to be associated with a stronger BOLD effect (see S1 Text). Both experiments converged to show that F2 variation was specifically tracked by neural activity in the right posterior superior temporal gyrus (pSTG), while perceptual decisional effort involved several regions of bilateral inferior prefrontal and posterior temporo/parietal cortex, and the right anterior temporal pole (Fig 1C, 2B). These activations, in particular the acoustic encoding of F2 variations, remained focal even at a lenient statistical threshold (S1 Fig). The spatial selectivity of the acoustic tracking of F2 was confirmed by a second fMRI study in which participants had to decide whether they heard /da/ or/ta/. In this case the morphed acoustic cue was no longer spectral (F2) but temporal (voice onset time). We found that this acoustic cue was encoded in a restricted region of the *left* STG and superior temporal sulcus (STS) (S8 Fig and S1 Text). In short, the right pSTG was recruited for encoding the slope of the 2^nd^ formant in the ba-da continuum, and the left STG/STS for encoding the duration of the consonant part in the data continuum, reflecting the hemispheric dominance for temporal vs. spectral acoustic processing (21).

**Fig 1.**
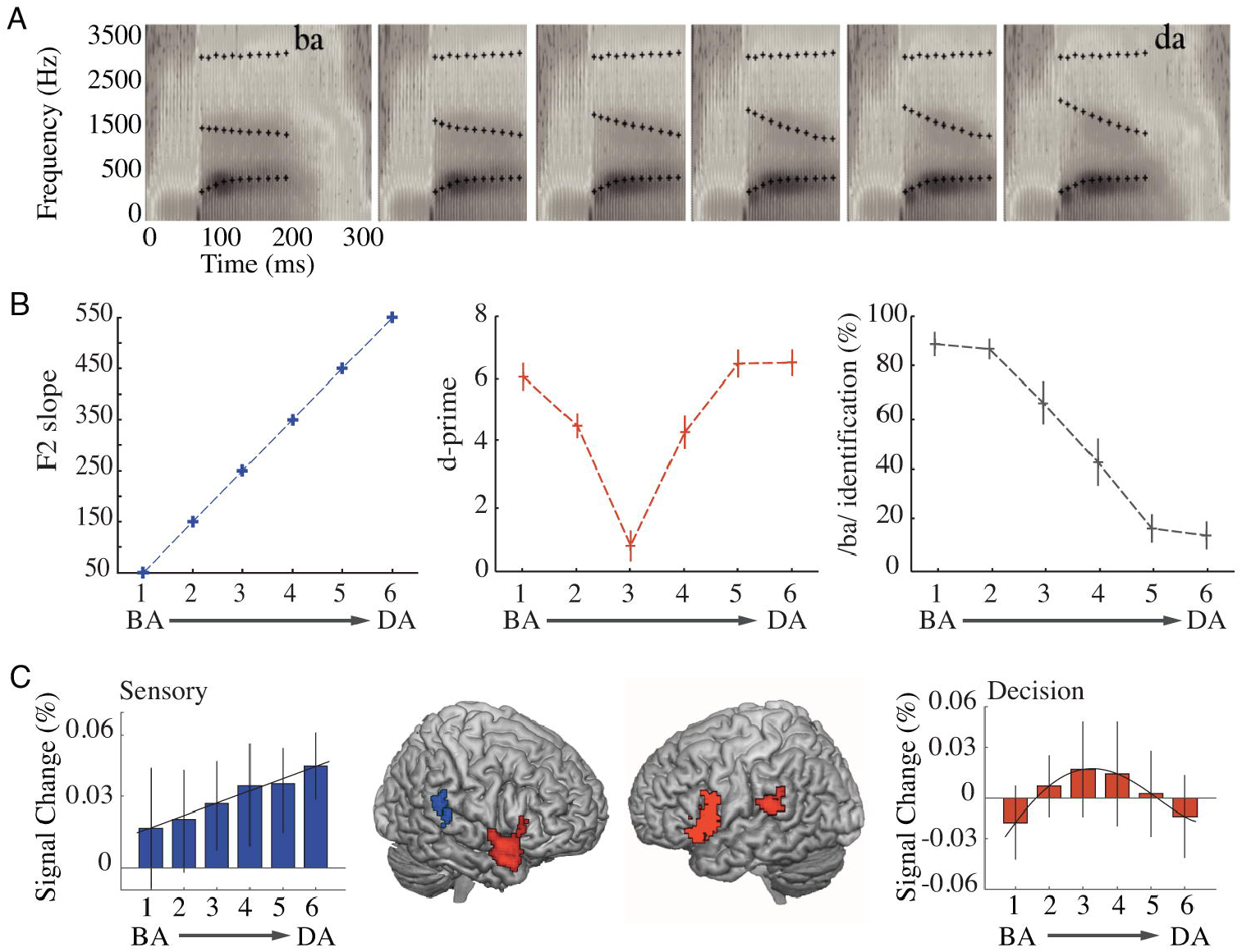
fMRI results. ***(A).*** *Spectrograms of the stimulus continuum between syllables /ba/ and /da/, synthesized with linear increase in F2-parameters (1650:100:2150 Hz). Full spectrograms at the extremes of the continuum represent /ba/ and /da/ prototype syllables (left and right panels, respectively). Middle spectrograms are centered on the F2-parameters.* ***(B)*** *Values for F2-parameters (in blue, left panel), average d-prime (in red, middle panel), and percent syllables identified as /ba/ (in grey, right panel) (mean* ± *s.e.m.) (C). Results of the regression analysis. Top panel: spatial localization of F2-parameters neural encoding (in blue) and d-prime (in red) in fMRI BOLD signal, expressed as beta coefficients. Significant clusters were found in the right posterior superior temporal lobe (pSTG)* (peak MNI coordinates, *x, y, z = 42, −34, 7, T = 3.21) for the F2-tracking, and in left posterior temporo-parietal (x, y, z = −51, −28,16, T = 4.41) and bilateral inferior prefrontal (x, y,z = 45,17, −5, T = 5.26; x,y,z = −48, 8, 22, T = 5.29) cortices for auditory perceptual decision (d-prime). Images are presented at a whole-brain threshold of P<0.01 (uncorrected). Bottom panel: percent signal change in the left inferior prefrontal cortex (in red) and in the right pSTG (in blue). BOLD signal increases with F2-parameters in the right pSTG, and with auditory perceptual decision load in the left inferior prefrontal region (in red).*

We used dynamic source modeling of the MEG data to explore the dynamics of acoustic encoding and perceptual decision. We found neural correlates of F2 parameters encoding 120ms post stimulus onset in the right pSTG. Auditory perceptual decision-related activity appeared in this region at 165ms and co-occurred with a second peak of F2 encoding activity at 175ms (Fig 2B). In addition to the spectral response profile within the right pSTG and left prefrontal cortex, a Granger causality analysis across the two areas showed that neural activity related to F2 and negative d′ (–d′) corresponded to bottom-up encoding and top-down decoding activity, respectively. Both analyses were associated with neural activity in the highgamma band for F2 variation and in the beta band for–d′, confirming the generic implication of these two frequency ranges in bottom-up and top-down processes (22–24) (Fig 2C). The MEG findings thus support the straightforward scenario in which auditory decisions arise from a focal readout of the region that encodes the critical sensory cue (F2) by prefrontal regions (25–27).

**Fig 2.**
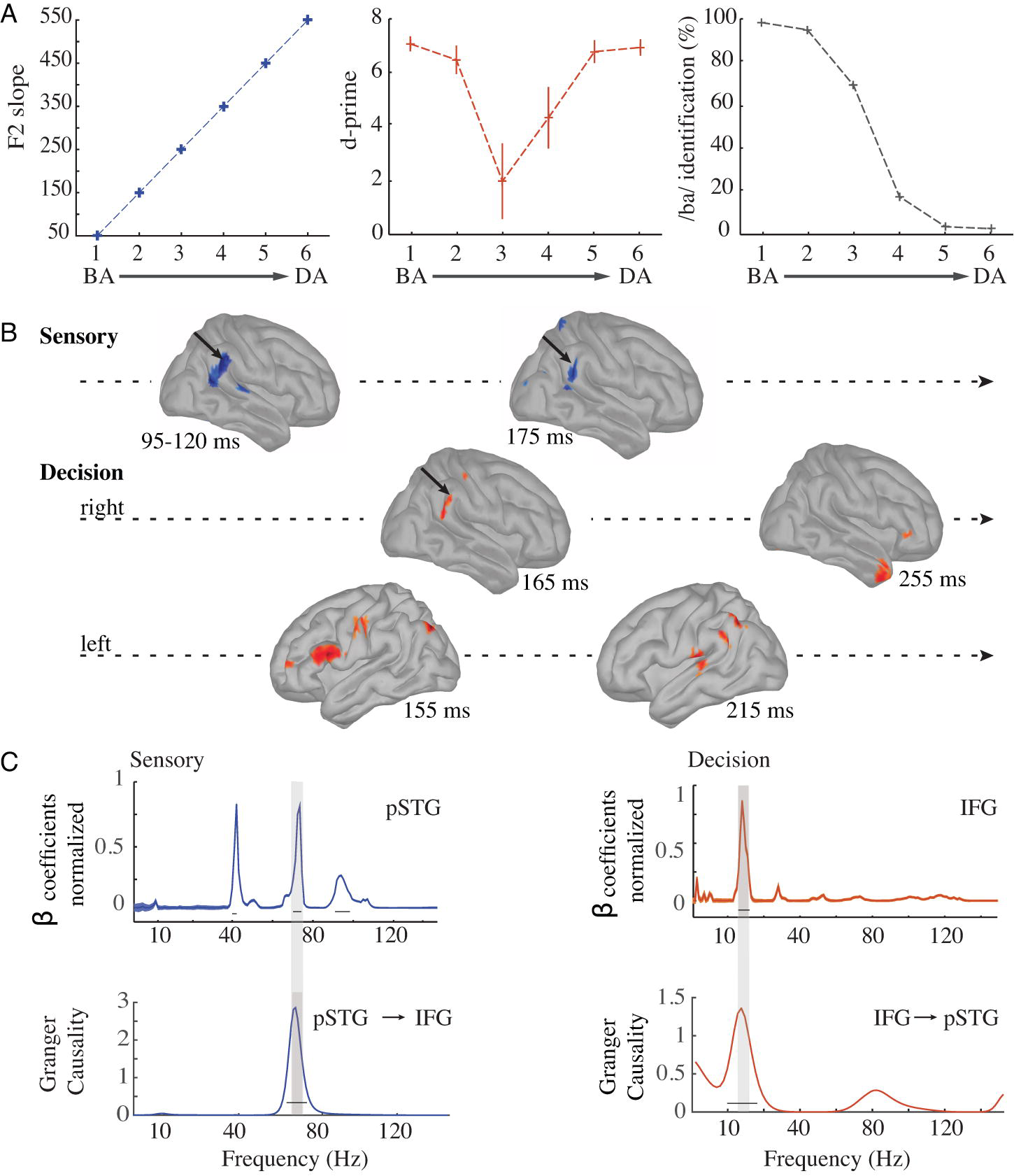
MEG results. ***(A).*** *Values for F2-parameters (in blue, left panel), average d-prime (in red, middle panel), and percent syllables identified as /ba/ (in grey, right panel) (mean* ± *s.e.m.).* ***(B).*** *Dynamic spatial localization of the neural encoding of F2 (in blue) and d-prime (in red) in MEG signals, expressed as beta coefficients. Only the bootstrapped P=0.05 significance threshold (Bonferroni-corrected) activations are represented. The right pSTG (pinpointed with black arrows) is first activated from 95 to 120ms for F2-parameters encoding, and then reactivated for phonemic decision around 165ms. (C). Top panel: Spectral profile of beta coefficients from regressions between F2 values and neural response in the right pSTG (left panel), and between –d’values and neural response in the left inferior prefrontal area (right panel). F2 was dominantly tracked by gamma and high-gamma activity, whereas decisional activity was expressed in the low beta band. Thick black lines indicate significant beta coefficients at P<0.05 (Bonferroni-corrected). Bottom panel: Granger Causality results between right pSTG and left inferior frontal gyrus (IFG). Thick black lines indicate significant granger coefficients at P<0.05 (Bonferroni-corrected). Top and bottom panels: shaded grey areas highlight the correspondence between beta coefficients and Granger Causality peaks. Left panel: high-gamma band for bottom-up activity from right pSTG to left IFG. Right panel: beta band for top-down activity from left IFG to right pSTG.*

Having established the global representational validity of the regions encoding the sensory features of interest (F2 variations), we then sought to examine the responses of these regions at a finer-grained scale using invasive electrophysiology. To maximize signal-to-noise ratio and spatial specificity in the exploration of coincident neural responses to F2 and auditory perceptual decision, we acquired intracortical EEG (i-EEG) data in 3 epileptic patients who together had 14 electrode shafts throughout the right temporal lobe (70 contact electrodes). Among them, one electrode shaft penetrated the right temporal cortex through Heschl’s gyrus (see Fig 3B). The deepest contacts of this auditory shaft strictly colocalized with the region that fMRI detected for F2 variation tracking. The patients performed the same syllable categorization experiment on a ba/da/ga continuum where the only changing acoustic cue was the F2 parameters (Fig 3A). Behavioral results show a good detection of ba and da, and a slightly less frequent detection of ga (Fig 3C). Strong evoked responses to syllables were only present in the auditory shaft, and were more marked/consistent in its two deepest contacts (Fig 3D, top row); the responses were weak to inexistent elsewhere (Fig 4A, colored plots). Significant F2 tracking was consistently detected in all auditory contacts (see Methods), with strong and structured effects in the two deepest ones (Fig 4D, middle row). Fully consistent with the MEG results, F2 values were encoded by broadband gamma activity (40-110 Hz) from about 150ms post-stimulus onset onward, i.e. 50ms after F2 appeared in the acoustic signal. Structured and strong neural activity related to F2 tracking was not observed in any of the other contacts of the same patient (Patient 1, S2 Fig). These data confirm the extreme spatial selectivity of F2 parameters encoding, and show that the encoding parameters of the discriminant acoustic cue are available in the pSTG for syllable recognition.

**Fig 3.**
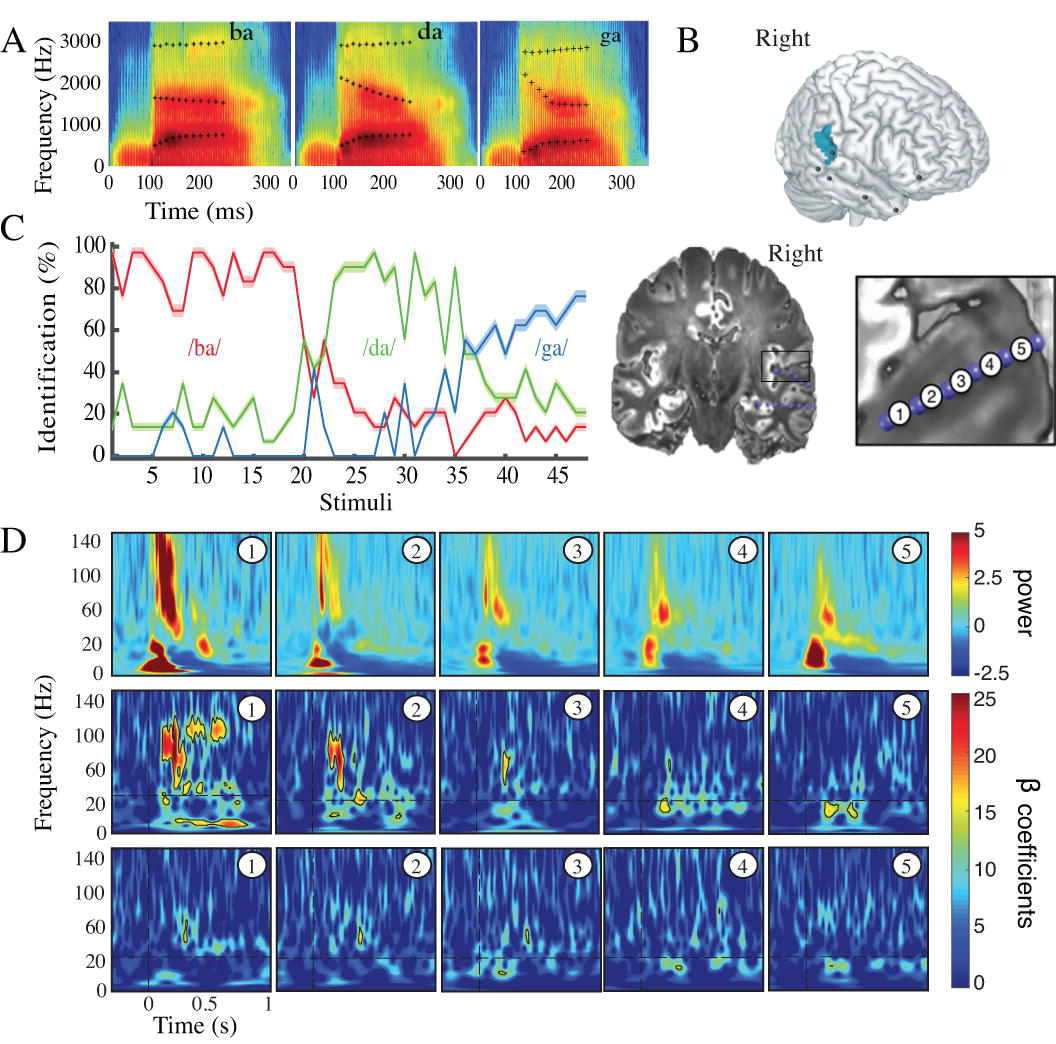
i-EEG results in Patient 1. ***(A).*** *Spectrograms of /ba/, /da/ and /ga/ prototype stimuli, synthesized with linear parametric F2-parameters.* ***(B).*** *Locations of i-EEG contacts in Patient 1. The auditory shaft labeled shaft 1 penetrated the right pSTG and Heschl’s gyrus. The patient had 5 other shafts distributed in the right temporal lobe. Bipolar montages from adjacent contacts are shown in the bottom right figure.* ***(C).*** *Percentage of syllable identification for each category for the three patients. The shaded zone indicates SEM. The first 20 stimuli are categorized as /ba/, the next 13 stimuli are categorized as /da/, and the last 11 stimuli are categorized as /ga/.* ***(D).*** *Top panel: evoked activity, averaged across stimuli, on each bipolar montage from the deepest (number 1) to the external-most (number 5) contact in the auditory shaft (shaft 1). Middle panel: time-frequency representations of beta coefficients from regression of F2 values against evoked activity on each contact of the auditory shaft. Significant F2 tracking was found in all contacts of the auditory shaft, with stronger effects in the two deepest contacts. Bottom panel: time-frequency representations of beta coefficients from regression of d-prime values against evoked activity on each contact of the auditory shaft. Decisional effects were significant on the third auditory contact about 200ms post-stimulus onset in the beta band. Middle and bottom panels: The vertical dashed lines indicate stimulus onset. The horizontal dashed lines indicate a change in the scaling of the oscillatory power for each time point and each frequency, with a 0.5 Hz resolution below 20 Hz and a1 Hz resolution above. Black contours indicate significant t-tests at q<0.05 (FDR correction).*

Decisional effects were globally weak in i-EEG signals, yet significant in the third auditory contact about 200ms post-stimulus onset in the beta band, in agreement with the MEG results, and in the deepest auditory contact about 350ms post-stimulus onset in the gamma band (Fig 3D, bottom row). Because both fMRI and MEG showed correlates of decisional effort at several other locations in the frontal and temporal lobe, we broadened the analysis in this patient to all contacts of each shaft (S3 Fig). Perceptual decision-related effects were weak, sporadic and inconsistent. They were significant before 500ms post-stimulus at only two other locations outside Heschl’s gyrus: in the right inferior prefrontal cortex (shaft 6, contact 1; consistent with fMRI, Fig 1C) and in the anterior temporal lobe (shaft 4, contact 4; consistent with MEG, Fig 2B).

We then sought to address whether focal neural activity could afford syllable categorization. Contrary to previous findings based on ECoG signals (1,2), local evoked activity from one contact was sufficiently discriminable to permit syllable categorization using a machine-learning algorithm (maximum correlation coefficient classifier, see methods). Decoding was possible from all individual auditory contacts, but worked best from the deepest one (Fig 4A, bar plots, S6A Fig). Within the other electrode shafts, univariate decoding based on single contact information was never possible. However, significant multivariate decoding from pooling all contacts in each shaft was significant for shafts 1, 2, 3, 4 and 6, even though it included non-responsive contacts. Reciprocally, multivariate decoding was not possible in the temporal pole shaft (shaft 5), even though we detected significant perceptual decision-related neural activity in this region with fMRI and MEG.

We subsequently addressed the key question whether the information used by the classifier corresponded to that used in the human decisional process. We examined whether there was a temporal correspondence between the dynamics of decoding, as assessed by time-resolved classification (28,29), and the presence of time-frequency neural cues that informed the subject’s perceptual decision. For this analysis, to ensure the independence of the analyzed dataset (S1 text), we no longer probed the decisional effort (search for information, –d′), but the decisional outcome. We approximated the neural cues that were critical to the decisional outcome by the difference in the time-frequency response between correctly and incorrectly recognized prototype syllables. The correct-incorrect contrast indicates the parts of the neural signal, which, if missing, are associated with an erroneous perceptual decision. Note that this contrast matches, as closely as possible, the output of the maximum-correlation coefficient classifier, which tests the extent to which a linear association can correctly predict syllables from neural activity.

**Fig 4.**
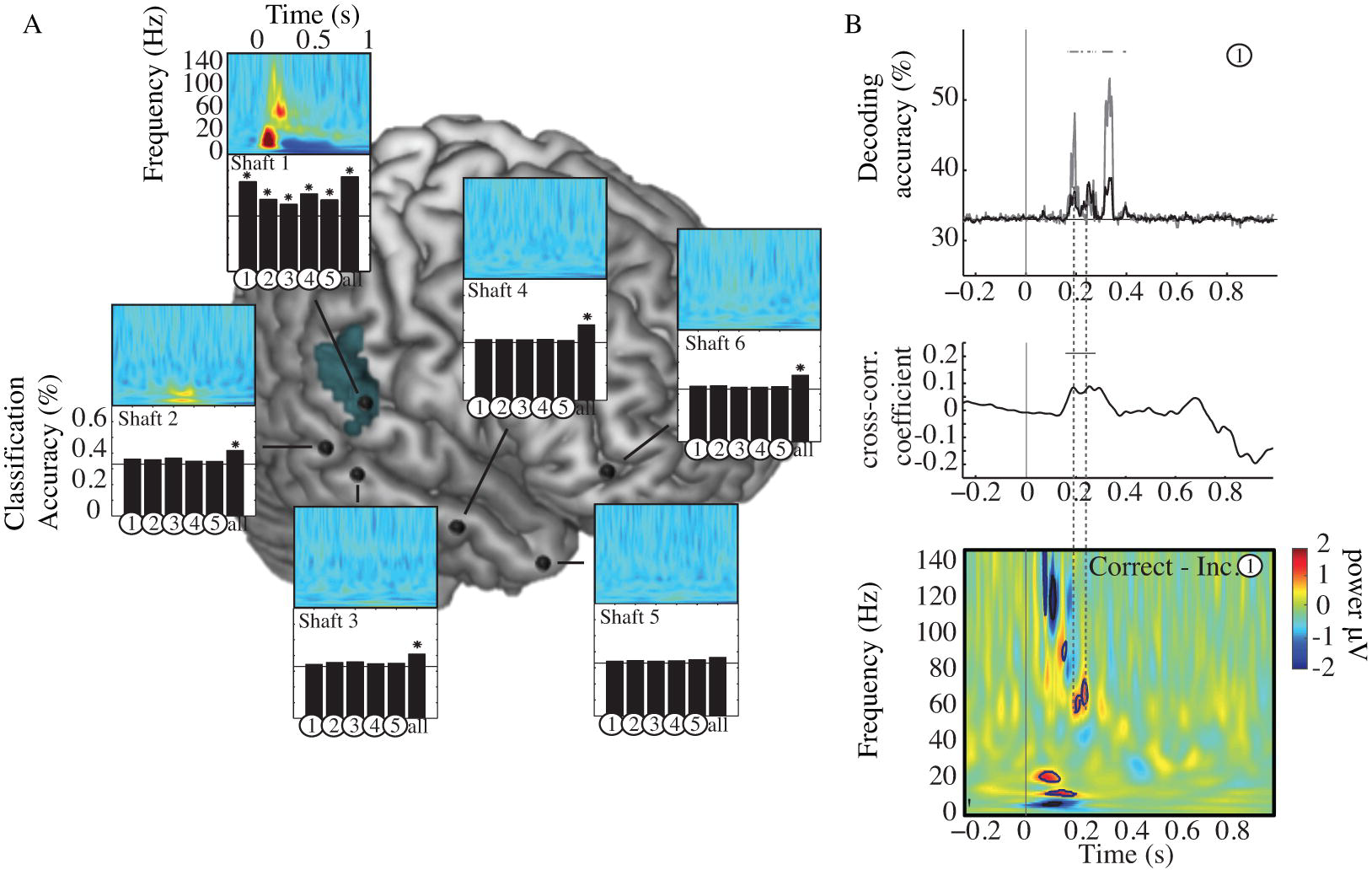
Decoding in Patient 1. *(A). Top graphs: time-frequency representation of evoked activity on each shaft. A strong evoked response to syllables is only present in the auditory shaft. Bottom graphs: neural decoding through univariate and multivariate classifiers. Histogram bars numbered from 1 to 5 show the univariate classifier results based on the activity from each contact of each shaft; the right bar (“all”) shows the multivariate classifier results based on multidimensional information from all contacts of each shaft. Stars above the black bars signal significant classification accuracy for specific contacts within each electrode shaft (q<.05, FDR-corrected). Univariate classification was possible from all auditory contacts (shaft 1) overlapping fMRI F2-parameters activation (blue shaded area), but worked best in the deepest one. Univariate classification failed everywhere except for these auditory contacts, whereas syllable decoding worked above chance using the multivariate approach in shafts 1, 2, 3, 4 and 6. (B). Top panel: Temporal relationship between the time-course of machine-decoding from the deepest auditory contact (upper panel, black line), mean univariate classification from all auditory contacts (upper panel, grey line) and the time-frequency cues used by the subject to make a correct perceptual decision (difference in the time-frequency response between correctly and incorrectly recognized syllables, lower panel). The grey thick lines show significant results for each time point (significant decoding accuracy, q<.05, FDR-corrected). Middle panel: cross-correlation coefficients between univariate decoding accuracy and significant correct-incorrect time/frequency clusters. Significant effects were found in the 60-80 Hz high-gamma band. The horizontal black line indicates significant cross-correlation coefficients at P<0.05 (Bonferroni-corrected). Bottom panel: Time-frequency differences between correct minus incorrect classification computed on contact 1 of shaft 1. Black borders indicate significant differences in neural activity between correct and incorrect classification scores, in comparison to a zero-mean normal distribution at q<.05, FDR-corrected.*

Significant time-frequency correlates of correct classification were only found in the three deepest contacts of the auditory cortical shaft (S4 Fig); they were sporadic and inconsistent elsewhere (red frames in Figure S4 show significant activity for t<500ms). In the deepest auditory contact (contact 1 on shaft 1) where both F2 tracking and univariate *classification* were maximal (Fig 3 and S5 Fig), cues associated with correct perceptual decisions were present as early as 150ms, i.e. before the first significant decoding peak (200ms) (Fig 4B, S4 Fig and S6B Fig). This important finding shows that within 150ms, the right pSTG had encoded enough information about F2 onset frequency and slope to inform human correct syllable recognition, and that this information could be exploited by the classifier to distinguish across syllables (see discussion).

**Fig 5.**
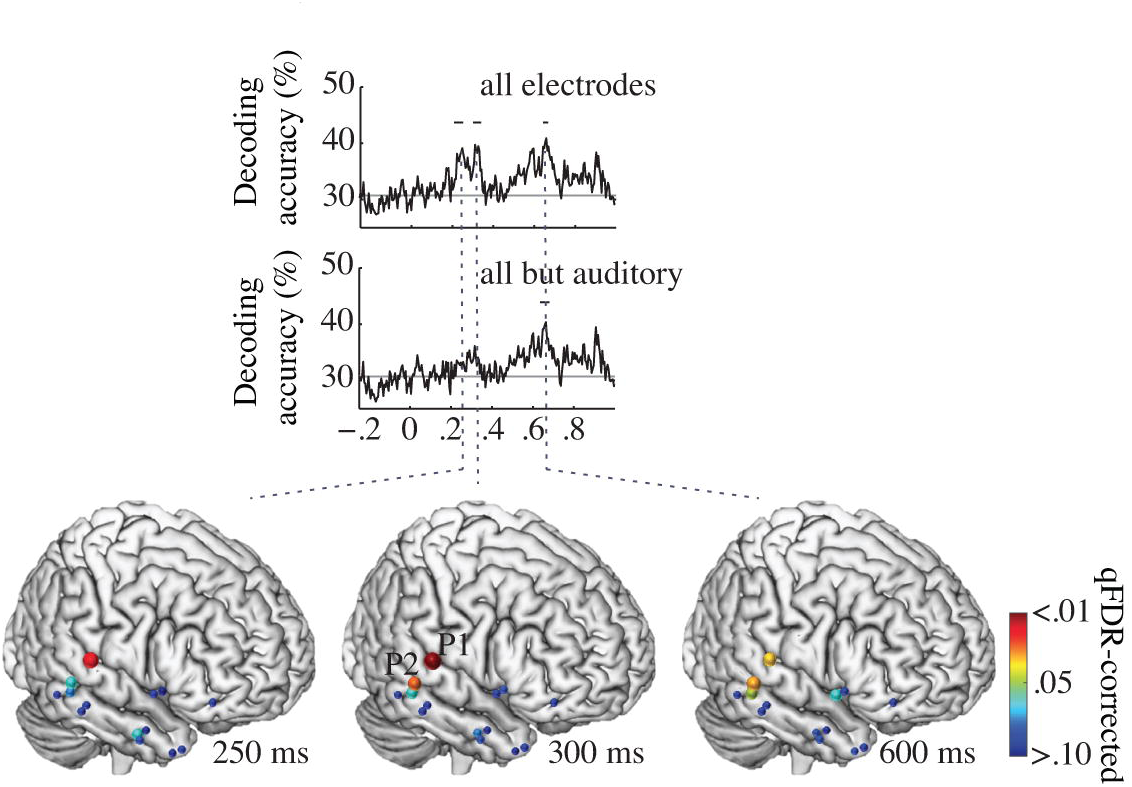
Decoding in all patients. *Time-course of the decoding accuracy from multivariate pattern classification with all shafts (upper panel), and without the auditory shaft of Patient 1 (lower panel). Early classification dropped below statistical significance, while the latest classification peak at 600ms remained unaffected. Location of shafts (n = 14) from which neural activity was recorded during syllable identification (3 patients, fixed-effects model). Colored dots show cluster-level significance from q>0.10 to q<0.01 (q-FDR corrected) multivariate classification (dot size proportional to q) performed on all shafts. Significant classification was observed at 250, 300 and 600ms, showing that syllables could be decoded from broadly distributed activity.*

Critically, the temporal coincidence between neural correlates of response correctness and machine-decoding (Fig 4B) was only partial. It was fairly good for the first two significant decoding peaks (<200ms) of both single auditory contact decoding and mean univariate decoding across all auditory contacts, but poor for the latest (and strongest) peak. The first two decoding peaks precisely coincided with transient high-gamma activity on the 60-80 Hz range (significant zero-lag cross-correlation, Fig 4B), in line with the observation made with MEG (Fig 2) that F2 was specifically encoded by neural activity in this frequency range, and that 60-80 Hz activity preceded decisional activity in the left IFG. However, the third decoding peak had no matching time-frequency event in the correct vs. incorrect activity profile (Fig 4B). These observations indicate that the classifier did not systematically capture those neural cues that informed the subject’s decision. Thus, machine-classification and human subjects did not exploit the same cues locally. Presumably the outcome of the mean univariate classifier reflected distributed information that was no longer relevant for – or assimilated into – neuronal decision variables. The strongest local decoding peak occurred at 370ms, i.e., more than 250ms later than the first correctness effect, about 100ms after the last one, and likely reflected post-decisional choice-correlated neural activity.

So far, the results indicate that phonemic decision was informed by focal early neural activity (<200ms) that encodes F2 in sustained multi-frequency oscillatory activity (Fig 2). Yet, distributed subthreshold neural activity, not detected by conventional (univariate) analyses of fMRI, MEG and intracortical EEG data, might also contribute to syllable identity encoding. We therefore addressed whether decoding was possible, even from contacts where there was no detectable F2 and from perceptual decision-related activity (S2 Fig, S3 Fig). We broadened the analysis to the 14 shafts of the 3 patients, including two more patients who had electrode shafts over the right temporal lobes (n=14), and performed time-resolved multivariate decoding from all cortical contacts (n=36). Significant decoding was found at 250, 300 and 600ms showing that syllables could be decoded from broadly distributed activity (Fig 5). To address whether distributed activity was driven by local auditory activity, we performed the same analysis without the contribution of the auditory shaft of Patient 1. Early classification (<300ms) dropped below statistical significance, but the latest classification peak at 600ms remained unaffected (Fig 5). This result demonstrates that decoding remained possible from cortical contacts that showed neither F2 nor auditory perceptual decision-related activity. We even obtained significant late decoding when deep structures, such as the amygdala and the hippocampus, were included in the multivariate analysis (n=70 contacts). As each penetrating shaft, except the auditory one, spanned functionally different territories, from the cortex to deeper structures, these findings show that the possibility of decoding neural activity in a multivariate approach does not allow one to conclude that the regions sampled for decoding amount to a meaningful neuronal representation, defined operationally in terms of a correct perceptual categorization. Overall, classification analyses from the i-EEG data confirmed the spatial selectivity of the early critical information involved in ba/da/ga syllable categorization. They also showed that syllable classification was possible from distributed activity (Fig 4 and Fig 5) that occurred later than the perceptual decision-related effects, as detected with both MEG and i-EEG.

Since the decoding of i-EEG returned positive results when pooling together contacts where no significant neural activity could be detected, we sought to explore the spatial distribution of /ba/ and /da/ category decoding using the MEG dataset. The idea was to determine whether whole brain decoding would be restricted to regions that showed statistically significant activation with all three approaches (MEG, Fig 2; fMRI, Fig 1; and i-EEG, S2 Fig and S3 Fig), or also to regions that did not critically participate in the task. This analysis was expected to provide time-resolved information to appraise whether decoding reflects non-critical processes downstream to sensory encoding and early decisional steps. Such a finding would concur with the i-EEG to suggest that decoding is possible from brain regions that are only collaterally involved in the cognitive process at stake.

Using a whole-brain sensor-based time-resolved multivariate learning algorithm (maximum correlation coefficient classifier, see Methods) (30), we found that speech sound categories could be decoded from very early brain responses in a focal region of the right pSTG (Fig 6). When we focused our analysis to those sensors that contained significant information about syllable identity (see Material and Methods), we found that activity recorded by the sensor MAG 1331 could be categorized as early as 100ms, with up to 78% accuracy (t = 13.01, q < .001, Cohen’s d = 4.7). Critically, syllable identity could also be decoded at a later time point, 220ms, on MAG 2011 first, and then at 350ms on MAG 0211 with scores reaching respectively 63% accuracy (t = 7.17, q < .001, Cohen’s d = 2.5) and 58% accuracy (t = 6.37, p < .001, Cohen’s d = 2.2). Corresponding source analyses then revealed that the decoding at 220ms post-stimulus onset arose from left STG and STS, two regions that were not primarily responsive to acoustic cue tracking and perceptual decision (Fig 2). Source analysis further showed that the decoding at 350ms arose from the left IFG, thus corresponded to late decisional effects. Together, these results show that, while decoding is most accurate in the region that critically encodes the acoustic information (the right pSTG), it is also subsequently possible from noisier activity in a broad left-lateralized network that contains associative information about the selected syllable. Interestingly, significant decoding was seen again in the right pSTG, 500ms post-stimulus onset with very high accuracy (89%, t = 15.10, q < .001, Cohen’s d = 5.5). This suggests that information propagation occurring across the whole language network between 100 and 500ms contributed to improve the quality of the categorical representations at a post-decisional stage.

**Fig 6.**
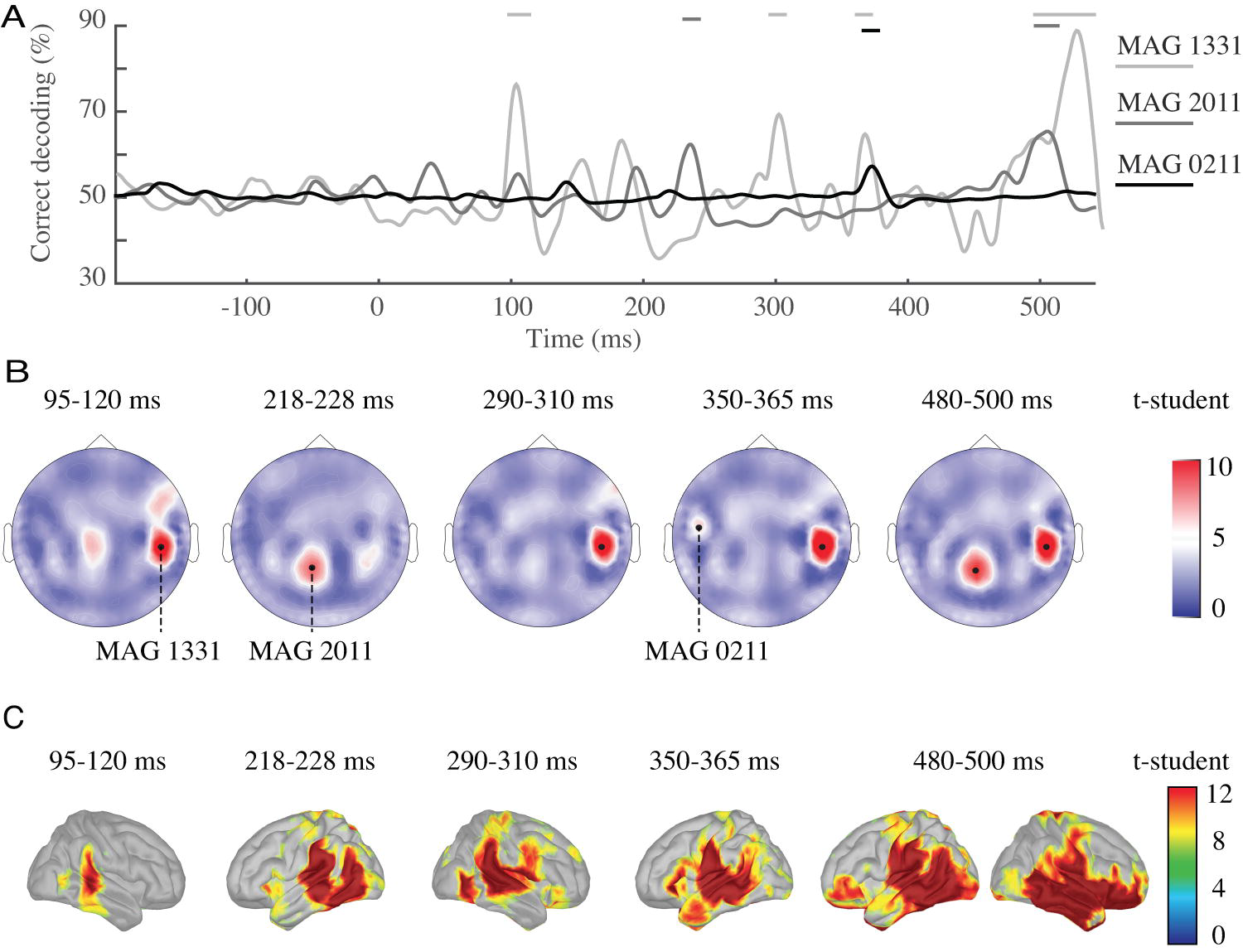
Decoding of MEG data reveals bilateral tempo-frontal cortex involvement in speech sound categorization. ***(A).*** *Percentage of correct decoding over time, for each of the magnetometers (MAG 1331, MAG 2011 and MAG 0211) showing significant decoding activity. X-axis: zero corresponds to stimulus onset; y-axis: 50% indicates chance performance. Horizontal dark and light grey lines indicate significant decoding (q < .05, FDR-corrected).* ***(B).*** *Sensor topographies depicting the average decoding response in magnetometers averaged within each of the five windows of interest. Black dots indicate the position of the 3 magnetometers showing significant decoding activity. (C). Source localization at key decoding times. Source localization for MEG signals displayed on a standard cortex at 100ms, 220ms, 300ms, 360ms and 500ms poststimulus onset. Right color bars indicate t-values.*

## Discussion

In this series of studies, we addressed whether human listeners’ perceptual decisions about the identity of speech sounds were grounded on a parallel readout of distributed representations (31), or whether, following more closely the principles of hierarchical processing, syllable identification in a categorical choice was informed by the efficient readout of a restricted brain area that contains limited but key neural information, as recently shown in monkeys (20). The current data converge to show that correct decisions about speech sound identity were informed by local and time-limited information (<300ms) present exclusively in the right pSTG.

Even though phonemic contrasts are most often associated with activations of the *left* STG (32), the *right* STG performs low-level spectral-based acoustic processing that is relevant for speech decoding (33,34) and categorization (35,36). Note that the right specificity of speech-related operations is easily missed when acoustic processing is not explicitly orthogonalized from decision-related neural activity (37). Our findings confirm that spectral- and time-based analyses of speech sounds involve the *right* and the *left* STG, respectively (see Fig 1 and S8 Fig). Our findings also confirm that in acoustically challenging situations such as those used in the current experimental design, the *left* IFG is mobilized (38) and interacts with the temporal cortex in sequential loops of bottom-up and top-down processes (37,39–41). Importantly, our fMRI and MEG results conjointly show that the focal readout of sensory encoding regions by prefrontal regions accounts for the decision variability relative to the perceptual state, and that the magnitude of neural activity associated with sensory processes depends on the discrepancy between heard and expected signals (42,43) (see Fig1C and Fig 2C).

Whether stimulus identity is retrieved from focal neural activity or from distributed information is a fundamental issue in neural coding theory. Our MEG and iEEG decoding results both show that the right pSTG is critical for perceptual decisions while distributed activations across frontal and temporal cortices reflect the reuse of sensory information for higher-level operations, such as meaning extraction, audio-visual integration, etc.. It has repeatedly been observed that behavior is as well explained by taking the activity of one or a limited set of neurons as it is by considering a large-scale population (44,45). This puzzling observation is backed-up by computational models showing that pooling together information from more neurons does not improve behavioral sensitivity. This could reflect the fact that neuronal responses are strongly correlated both intrinsically by neural noise and by stimuli, so that the information contained in a neural population saturates as the number of neurons increases (46,47). Recently, a combination of experimental and theoretical data suggests that in most cases, behavior is best explained by optimal readout of a limited set of neurons (48). The authors then conclude that the neural code is redundant, and that some areas are decoded near-optimally, while others are not efficiently read out. They propose that deactivating these regions should not impact behavior.

Reciprocally, as we tested in a brain-damaged patient carefully selected with respect to the lesion extend and localization (S1 Text, S7 Fig), deactivating regions that are optimally decoded would be expected to impair behavior. The lesion of one of the two focal regions that we identified as key for F2-based syllable categorization dramatically disrupted performance. The impairment was selective to the type of stimulus but not to the type of task, as categorization of similar syllables (da/ta) based on temporal cues, i.e. VOT, was spared. This also was expected as syllable categorization based on VOT specifically involved the left middle temporal cortex (S8 Fig). The selectivity of the impairment with respect to the acoustic material shows that for this type of task there was no distributed rescue system. These behavioral data in a single patient remain of course non generalizable and further lesion data will be necessary to confirm our results. Moreover, a full demonstration that, even though phoneme categories are represented in a distributed fashion, focal/early sensory information is predominantly used to categorize speech sounds, would strictly speaking require showing that distributed lesions do not impair task performance. This demonstration, however, is impossible to make because at the extreme, if the whole language system is injured, no linguistic operation can be achieved at all. Although using a variety of approaches we tried to collect as many arguments as possible to address how information is used for making perceptual decisions about speech sounds, we acknowledge that the present study can only be partly conclusive.

Critically, however, our results show that there is a fundamental distinction to be made between the decoding of neuronal activity by machine-learning and neuronal decoding made by the brain to achieve a specific cognitive goal. We found that distributed noise-level neural information, which did not carry reproducible and statistically detectable information about the sensory cue (F2-parameters) or about the categorical decision process (49), was nonetheless sufficient to inform a classifier. While this confirms previous observations that phonemic information is present in a distributed manner (1,2), the fact that classification was possible only on late responses once the region encoding the critical acoustic cue (F2 slope) was removed from the multivariate analysis, suggests that distributed phonemic representations reflect the diffusion of the information throughout the language system. While distributed information might be useful for late association-type linguistic processes, it did not appear necessary for the categorization task, as correctness effects occurred focally and early in time. These findings show that neuronal activity containing information about speech sound categories is not uniformly *useful* for categorizing these sounds. More generally, our findings highlight the fact that decoding algorithms, even if they can make use of distributed information that might reflect a brain state context (45) or mental association, do not indicate which regions are necessary or causal for the cognitive process at stake – here perceptual decision-making. These results hence suggest that distributed information about categories might reflect the redundancy of noise-level information in the brain, i.e. associative neural activity, rather than a spatial neural code that is accessed in parallel when making a speech category perceptual decision.

Categorizing auditory syllables is a relatively low-level process, which could in theory be underpinned by distributed representations, and yet in our data this appears not to be the case. What then might be denoted by previously observed broadly distributed representation maps (8)? In particular phonemic maps organized along articulatory features (50)? Most of these important findings (1,40,51) were obtained in natural listening conditions. In Mesgarani et al. (1), for instance, maps were drawn from cortical surface neural activity measured 150ms after each phoneme in connected speech, implying that activity at each time-point integrated distributed activity of several preceding phonemes and reflected co-articulation and contextual associations from the preceding words. When passively listening to connected speech there was likely no explicit serial phoneme decoding, but rather a global readout of sentences, which likely required accessing several phonemic representations at once. In the same way as a computer keyboard spaces letters to minimize interference when typing them, our brain might organize the phonemic space as a function of how we need to retrieve them for production (42), i.e. following a feature-based logic. That this organization exists does not imply that it is “optimal” for perception, just as reading words through a keyboard spatial organization would largely be sub-optimal. The present findings confirm the existence of distributed phonemic representations, but also question the use our brain makes of them in an explicit speech perception context, as phoneme recognition does not seem to rely on distributed activity. Importantly however, it might be the case that during natural speech perception, cross-hierarchical readout of redundant/correlated neural activity is genuinely exploited as a form of trade-off between accuracy of single phoneme identification (focal) and joint access to multiple representations (distributed). It will be essential in the future to address whether sub-optimal decoding of large neuronal populations could be an optimal way to handle access to multiple representations.

## Materials and Methods

### Experimental Procedures

#### Subjects

Thirty-one healthy subjects participated in the MEG study (16 females – age range: 22-31 years), and 16 in the fMRI study (9 females – age range: 22–29 years). Intracortical EEG (i-EEG) was recorded in 3 epileptic patients (1 female – ages: 44, 25 and 65 years) who underwent surgery for clinical monitoring, and 1 patient (female – age: 77 years) was tested behaviorally 8 months after an ischemic stroke. All participants were right-handed, French-native speakers, and had no history of auditory or language disorders. The experimental protocols were approved of by the Inserm ethics committee in France (biomedical protocol C07-28) for the participation in the MEG and fMRI experiments, and by the University Hospital of Geneva in Switzerland (13–224) for studies in epileptic and stroke patients. All participants provided written informed consent prior to the experiment.

#### Stimulus synthesis and behavioral testing

High-quality speech stimuli were synthesized using STRAIGHT (52) together with an original morphing method, based on a modified LPC (Linear Predictive Coding) analysis-synthesis scheme (53). The second formant transition, was morphed in order to build a linear speech sound continuum between /ba/, /da/ and /ga/ (54,55). Through the continuum, we kept constant a low-pass filtered pulse train for the voiced part, a filtered noise throughout all the stimulus, an additional white noise for the burst, the global amplitude of the stimulus, and the first and third formant transitions. We also kept the same time dependency fo = f(t) for all stimuli, as extracted from the original ‘ba′. A 6 item /ba/ /da/ continuum was presented to healthy subjects and to the stroke patient. A longer 48 item continuum /ba/ /da/ /ga/ was used for testing epileptic patients in order to obtain more responses around syllable boundaries to compare correct and incorrect categorization. Note that /ba/ and /da/ categories only differed on the F2 dimensions, and hence processing this single cue was sufficient for correct perception. Another 6 stimuli /da/ /ta/ continuum was used for both the behavioral and second fMRI control experiments. In that continuum, we varied the length of the voice onset time (VOT) by deleting or adding one or several voiced periods in the signal, before or after the burst.

#### Tasks design

Auditory stimuli were presented using Psychophysics-3 Toolbox and additional custom scripts written for Matlab (The Mathworks, Natick, Massachussetts, version 8.2.0.701). Sounds were presented binaurally at a sampling rate of 44100 Hz and at a comfortable hearing level individually set before the experiment via earphones. Prior to the experiment, each participant undertook a short session during which the minimum amplitude level leading to 100% categorization accuracy was estimated using an adaptive staircase procedure. This threshold was used to transmit the stimuli (mean 30 dB sensation level). Each continuum was delivered to participants in two independent sessions of 240 trials each for fMRI recording and of 270 trials each for MEG recording. 144 trials constituted the experiment used for epileptic patients.

Participants were asked to perform an identification task. Each trial comprised one sound (randomly chosen among the 6 or 48 stimuli of the continuum), followed by a 1s silence gap; then, a response screen with ‘ba′ and ‘da′ written syllables (in MEG, fMRI, and behavioral sessions) or ‘ba′, ‘da′ and ‘ga′ syllables (in i-EEG sessions) was displayed. Syllables were randomly displayed from right to left on the screen to prevent motor preparation and perseverative responses. During fMRI recording, the appearance of the response screen was randomly jittered 100, 300 or 500ms after the silence gap. Participants indicated their response on the syllable by pressing the corresponding left or right button response as quickly as possible. Subject’s responses were purposely delayed, and response times hence do not constitute relevant data. To prevent eye movements, subjects were asked to fixate the central cross and blink only after giving their motor response. After the response, a jittered delay varying from 3 to 5 s led to the next trial.

#### MEG recording and preprocessing

Continuous cerebral activity was recorded using an Elekta Systems MEG device, with 102 triple-sensor elements each composed of 2 planar gradiometers and 1 magnetometer. MEG signals were recorded at a sampling rate of 1 kHz and online bandpass filtered between 0.1 Hz and 300 Hz. A vertical electro-oculogram (EOG) was simultaneously recorded. Before MEG recording, headshape was acquired for each participant using Polhemus. After the MEG session, an individual anatomical MRI was recorded (Tim-Trio, Siemens; 9 min anatomical T1-weighted MP-RAGE, 176 slices, field of view = 256, voxel size = 1 × 1 × 1 mm^3^). MEG data were preprocessed, analyzed and visualized using dataHandler software (wiki.cenir.org/doku.php), the Brainstorm toolbox (56) and custom Matlab scripts. A Principal Component Analysis (PCA) was performed through singular-value decomposition function of numerical recipes, to correct artifacts (low derivation). The first two components from the PCA were zeroed and signal matrix was recomputed. PCA rotated the original data to new coordinates, making the data as flat as possible. The data were then epoched from is before syllable onset to 1s after syllable offset. Another PCA was then performed on the epoched data when blinks occurred. PCA components were visually inspected to reject the one capturing blink artifacts. On average, 2.1% (±0.7%) of trials per participant (mean ± SEM) were contaminated by eye movement artifacts and were corrected before further analyses.

#### fMRI recording and preprocessing

Images were collected using a Verio 3.0 T (Siemens) whole body and radio frequency coil scanner. The fMRI blood oxygenation level-dependent signal (BOLD) was measured using a T2*-weighted echoplanar sequence (repetition time = 2110 ms; echo time = 26 ms; flip angle = 90°). Forty contiguous slices (thickness = 3 mm; gap = 15%; matrix size = 64 × 64; voxel size = 3 × 3 × 3 mm^3^) were acquired per volume. A high-resolution T1-weighted anatomical image (repetition time = 2300 ms; echo time = 4.18 ms; T1 = 900ms; image matrix = 256 × 256; slab thickness = 176 mm; spatial resolution = 1 × 1 × 1 mm^3^) was collected for each participant after functional acquisition. Image preprocessing was performed using SPM8 (The Wellcome Trust Centre for Neuroimaging, University College London, UK, http://www.fi.ion.ucl.ac.uk/spm/). Each of the 4 scanning sessions contained 400 functional volumes. All functional volumes were realigned to the first one to correct for interscan movement. Functional and structural images were spatially preprocessed (realignment, normalization, smoothed with an 8-mm full-width-at-half-maximum isotropic Gaussian kernel) and temporally processed using a high-pass filter with a cutoff frequency of 60 Hz. We then checked data for electronic, and rapid-movements artifacts using the ArtRepair toolbox (http://cibsr.stanford.edu/tools/human-brain-project/artrepair-software.html). Artifacted volumes were substituted by linear interpolation between contiguous volumes and explicitly modeled in the following statistical analyses. Estimated head movements were small compared to voxel size (<1 mm), and 3.2% (±0.3%) of the volumes were excluded due to rapid head movements (>1.5mm/s).

#### Intracranial stereotactic electroencephalography (i-EEG) recording and preprocessing

Electrophysiological activity was recorded over arrays of depth electrodes surgically implanted to identify epilepsy focus. Intracranial EEG was recorded (Ceegraph XL, Biologic System Corps.) using electrode arrays with 8 stainless contacts each (AD-Tech, electrode diameter = 3 mm, inter-contact spacing = 10 mm), implanted in several brain regions in the right hemisphere (see Fig 3, 4, 5). We determined the precise electrode shaft locations by co-registering a post-operative computed tomography scan (CT) with a high-resolution anatomical MRI template. For the i-EEG recordings, we used bipolar montage where each channel was referenced to its adjacent neighbour. We sampled the i-EEG signal at 1024 Hz for patient 2, and at 2048 Hz for patients 1 and 3.

Steady-state frequency spectra were estimated using a standard Fourier transform from 1 s before the onset of the stimulus to 1 s after the offset of the stimulus. Time-frequency power was defined as the single-trial square amplitude estimates of complex Fourier components. Time-frequency analyses were carried out using the Fieldtrip toolbox for MATLAB (57). The spectral-power of MEG oscillations was estimated using a family of complex Morlet wavelets, resulting in an estimate of power at each time point and each frequency. We restricted the analysis to frequencies between 2 and 150 Hz, spanning the whole range of relevant brain rhythms. Note that time-frequency transform uses frequency dependent wavelets (from 3 to 7 cycles per window), with decreasing time-windows with increasing frequency.

#### Neural encoding of parametric information

We regressed out single trials of MEG, fMRI and i-EEG signals against i) the acoustic dimension, corresponding to F2-parameters (the onset value of the second formant (F2) and the F2-slope linearly co-varied in six steps) or to voice onset time (the voicing length before and after the consonantic burst varied in six steps), and ii) the categorization difficulty dimension corresponding to the inverse of the discriminability index from signal detection theory (*–d’*). These two dimensions are naturally orthogonal (r = .02, p >.20). A general linear regression model was carried out separately for each dimension (sensory encoding and decisional effort) along the stimuli, and finally averaged across participants to produce a group-level grand average. That approach was adopted to disentangle the neural correlates of basic bottom-up perceptual processing indexing the tracking of the acoustic cue, from the neural correlates of the categorization difficulty reflecting the distance of each stimulus to the phoneme identity criterion (37,58).

#### fMRI – Neural encoding of parametric information

Statistical parametric *t* scores were obtained from local fMRI signals using a linear multiple regression model with sensory encoding (F2-parameters or voice onset time value for each condition) and decisional effort (–*d*‘ value reported by each subject for each trial) as covariates. Regression parameters were estimated in every voxel for each subject, and parameter estimates were then entered in a between-subject random-effects analysis to obtain statistical parametric maps. We identified brain activation showing significant contrasts of parameter estimates with a voxelwise (T = 3.21, p < .005, uncorrected) and clusterwise (p < .05, uncorrected) significance threshold. Anatomical locations were determined based on automated anatomical labeling (AAL). Regressors of interest were constructed by convolving functions representing the events with the canonical hemodynamic response function. For each continuum, a categorical regressor modeled the ‘sound′ event using a Dirac function time locked to syllable onset. Two hierarchically orthogonalized parametric regressors (referred to as “sensory encoding” and “decisional effort” regressors) were added to the ‘sound′ regressor in order to capture the modulation of BOLD activity as a function of F2 variation tracking and categorization difficulty.

#### MEG – Neural encoding of parametric information

We first used single-trial signals on each sensor to perform parametric regressions at successive times from −0.2 to 1 second following stimulus onset. For each participant and each sensor, we calculated the time course of beta coefficients and then computed cortical current maps with Brainstorm using the weighted minimum-norm estimation approach – meaning that the time series for each source are a linear combination of all time series recorded by the sensors (59). Sources were estimated for each subject on the basis of individual MRI images. After realignment and deformation of each subject’s cortical surface, sources were projected onto the standard MNI/Colin27 brain, to perform grand mean averages. We then performed within group statistics so as to show the sensitivity to sensory encoding and decisional effort dimensions. Note that the two transformations applied to the data (regression and source-projection) both capture a *linear* relationship between the observed and the expected data, and can thus be implemented in whichever order. Nonetheless, we found that regressing data in the sensor space first, then “source projecting” the resulting beta values, was less sensitive to noise. Source projecting the results of the regression optimizes the signal-to-noise ratio (SNR) from the sensor data, and improves the interpretability of the source maps (Fig 2B). This method, therefore, was used to localize the sources of sensory and perceptual decision components.

Single-trial evoked signals on each sensor were also used to compute source current maps for each trial. The inverse operators were generated with the default MNE parameters and applied at the single-trial level. The estimated sources were morphed to the MNI brain. We then extracted single-trial neural activity from regions of interest defined according to the Destrieux atlas (60) (G_temp_sup-Plan_tempo, G_temp_sup-G_T_transv, G_front_inf-Opercular, G_front_inf-Triangul). Single-trial evoked responses projected on these selected sources were used in two ways:

(1) For each participant, we regressed out single-trial neural activity to estimate spectral power of the beta coefficients via a standard Fourier transform. Time frequency analyses were carried out according to the exact same parameters defined in the previous paragraph (*Intracranial stereotactic electroencephalography (i-EEG) recording and preprocessing).* We thus estimated the trial-to-trial variability in neural signal from region of interests at a given frequency that describe sensory encoding or decisional effort (t-test against zero, *P* < 0.05, *Bonferroni corrected*).
(2) For each participant, source activity in the pSTG and in the left inferior frontal gyrus (IFG) was used to measure Granger Causality (GC). While GC is classically used to assess causal influence between two time series, we here computed GC for non-stationary time series, such as oscillating neural signals (61,62). We used a non-parametric test by computing a spectral density matrix factorization technique on complex cross-spectra, obtained from the continuous wavelet transform of source-reconstructed MEG time series. We then assessed the linear directional influence between two brain areas, the pSTG and the left IFG.

GC was computed on a *trial-by-trial* basis for each subject (61), and averaged across time and trials. Because we computed GC on oscillating neural signals that are non-stationary in time, GC spectra were thus obtained in a non-parametric manner by computing Geweke’s frequency domain version of GC without going through the multivariate autoregressive model fitting. For each subject, we computed the mean GC across trials and the corresponding standard deviation. The original GC spectra were then standardized to obtain a vector of z-values, one for each frequency. Top-down and bottom-up influences were measured simultaneously, and information flow was considered primarily top-down when GC from the left IFG to right pSTG exceeded GC from pSTG to IFG, and bottom-up in the inverse case. We tested for significant frequency peaks separately for bottom-up and top-down GC direction, in directly comparing the z-transformed vectors obtained from GC spectra to a zero-mean normal distribution, and corrected for multiple comparisons with the Bonferroni method at p < .05. Our choice to focus on left-IFG was empirically motivated. Previous papers (e.g., (63–65)) have shown that the left-IFG is consistently involved in articulatory processing during speech perception but also in lexical information retrieval, both skills that are engaged when categorizing *ambiguous* speech sounds – i.e., when the internal perceptual decision criterion is difficult to reach, as in the current study.

#### MEG – Decoding analyses

Decoding analyses were performed with the Neural Decoding Toolbox (66), using a maximum correlation coefficient classifier on single-trial induced responses, across all MEG sensors. Data from both magnetometers and gradiometers were used. The analyses were performed with a cross-validation procedure where the classifier is trained on a subset of the data (80% of the data), and then classifier’s performance is evaluated on the held-out test data (20% of the data). Classification accuracy is reported as the percentage of correct trials classified in the test set averaged over all cross-validation splits. This procedure was repeated 50 times, leaving out each cross-validation split, and the final decoding accuracy reported is the average accuracy across the 50 decoding results. Additionally, an analysis of variance, based on second-level test across subjects, was applied to the test data to select at each time point those sensors that were significantly sensitive to syllable identity. We then assessed statistical significance using a permutation test. To perform this test, we generated a null distribution by running decoding procedure 200 times using data with randomly shuffled labels for each subject. Decoding results performing above all points in the null distribution for the corresponding time point were deemed significant with P < .005 (1/200). The first time decoding reached significantly above chance was defined when accuracy was significant for five consecutive time points. Source localization associated to the decoding results has been computed from evoked trials using the MNE source-modeling method (see above).

#### i-EEG – Neural encoding of parametric information

We performed the same parametric regressions on *i-EEG* recordings from Patient 1. These analyses have only been done on that patient, as he was the only patient for whom one shaft showed a significant induced response to syllable perception. Shaft 1 colocalized to the site where spectral cue tracking was found with fMRI and MEG (right posterior STG). We selected the 5 deepest contacts on each shaft; those contacts were located between the Heschl’s gyrus and the STG on shaft 1. Parametric regressions were carried out at successive times t from −0.2 to 1 second post-stimulus onset, on each selected bipolar derivation (i.e., from the deepest (1) to the most external (5) contact). We computed the power in each frequency band at each time point of each beta coefficient, with a millisecond resolution, similar to the induced power (between 2 and 150 Hz, with a 0.5 Hz resolution below 20 Hz and 1 Hz above, by applying a TF wavelet transform, using a family of complex Morlet wavelets; m = 3 to 7). For each contact, a null distribution was computed by repeating the identical regression procedure 1000 times with shuffled regressors. We used standard parametric tests (t-test against zero) to assess the statistical significance of each parametric regression. The type 1 error rate (False Detection Rate, FDR) arising from multiple comparisons was controlled for using nonparametric cluster-level statistics (67) computed across contacts, time samples and frequencies.

#### i-EEG – Decoding analyses

We used a maximum correlation coefficient classifier (Neural Decoding Toolbox (66)) on single-trial induced responses. We applied it on time series using a cross-validation procedure where the classifier is trained on 90% of the trials, and tested on the remaining 10%. Our recordings consisted in 3 repetitions of each stimulus condition (48 stimuli from /ba/ to /ga/) for Patient 1 and in 6 repetitions of each condition for Patients 2 and 3. This procedure was repeated 1000 times, and the results were averaged.

We estimated single-trial decoding of the neuronal response induced by different syllables using both uni- and multi-variate classification. The univariate classification was applied on each bipolar derivation (i.e., on each of the 5 contacts of each shaft) whereas the multivariate classification was performed on neural activity from every bipolar derivations of one shaft (i.e., on the 5 contacts of each shaft, pooled together), and then on all bipolar derivations of Patient 1 (on all contacts of the 6 shafts, pooled together), and finally on all three Patients (on all contacts of the 14 shafts, pooled together). Single shaft multivariate decoding was compared to mean univariate decoding computed first from each contact and then averaged. Decoding accuracy is expressed as % correctly classified trials in the test set. A null distribution was computed by repeating the identical classification procedure 1000 times with shuffled labels. We defined the number of classification repetitions with respect to the number of multiple comparisons done from each contact *(FDR-corrected for univariate decoding performed on each of the 5 contacts, time samples and frequencies; FDR-corrected for multivariate decoding performed on each of the 6 shafts, time samples and frequencies, FDR-corrected for multivariate decoding performed on all shafts together, time samples and frequencies).* Decoding accuracy was considered significant at q<0.05 if accuracy exceeded randomized classification at two consecutive time points.

#### *i-EEG – Differences Correct* minus *Incorrect.*

The psychometric identification function with percentage reporting /ba/, /da/ or /ga/ was defined along the corresponding continuum. Boundary separations determine the accuracy of categorical choice: the steeper the slope, the more accurate the perceptual decision. Patient’s ratings along the continuum were used to split responses into correct and incorrect trials. We subsequently computed the difference in neural activity from selected bipolar derivations between correct and incorrect conditions, and then compared it with the zero-mean normal distribution thresholding at q < 0.05 (*FDR*-corrected for multiple comparisons on shafts (30 shafts tested for patient 1), time and frequency dimensions). This procedure was repeated 1000 times with shuffled labels for correct and incorrect conditions.

## Author Contributions

S.B. and A-L.G. designed the study; S.B. and S.K. designed the stimuli; S.B performed the experiments; S.B., V.C., R.T., A-L.G. analyzed the data; S.B., V.C., A-L.G., D.VV. wrote the manuscript; A.G.G., M.S. recruited the patients. All authors contributed to the final version of the manuscript.

## Acknowledgements

We are grateful to Lorenzo Fontolan and Clio Coste for methods support, and to Aaron Schurger, Luc Arnal, Andreas Kleinschmidt, David Poeppel, Virginie van Wassenhove, Corrado Corradi Dell’Acqua, Narly Golestani, and Clio Coste for comments and useful discussions about earlier versions of this manuscript. Funded by ERC Compuslang GA260347 and SNF 320030_i493ig to A-L.G., by SNF 140332 and 146633 to M.S., by ANR-10-LABX-0087 IEC, ANR-10-IDEX-0001-02 PSL* (program “Investissements d’Avenir”), and ANR-16-CE37-0012-01 to V.C., and by SNF P300Pi_1675gi to S.B.

---

*S****1 Fig. fMRI ba/da experiment*** *Bilateral inferior prefrontal (red), left (red) and right (blue) pSTG, all displayed at a P=0.05 uncorrected threshold, k=20 (“any-depth” search mode).*

***S2 Fig. i-EEG results in Patient 1.*** *Time-frequency representations of beta coefficients from regression of F2 values against evoked activity on each contact of each shaft. Red frames indicate shaft contact where significant F2 tracking was found (comparison to zero-mean normal distribution thresholded at q < .05).*

***S3 Fig. i-EEG results in Patient 1.*** *Time-frequency representations of beta coefficients from regression of d-prime values against evoked activity on each contact of each shaft. Red frames indicate contacts where decisional effects were significant (comparison to zero-mean normal distribution, thresholded at q< .05).*

***S4 Fig. i-EEG results in Patient 1.*** *Difference in the time-frequency response between correctly and incorrectly recognized syllables for each contact of each shaft. Red frames indicate contacts where significant correctness effects were found (comparison to zero-mean normal distribution, thresholded at q < .05).*

***S5 Fig. Patient 1, decoding on auditory shaft (Shaft 1).*** *Significant univariate decoding was strongest on the 1^st^ contact, but even stronger using the multivariate approach.*

***S6 Fig.* Effect size (Cohen’s d). *(A).*** Effect size (Cohen’s d) of machine-decoding. For univariate decoding, contacts 1 and 2 of shaft 1 showed a *large* effect size (Cohen’d > .8), whereas contacts 3, 4 and 5 of shaft 1 showed a *medium* effect size (.5 < Cohen’s d < .8). A *small* effect size (Cohen’s d < .2) was found for all contacts of shafts 2, 3, 4, 5 and 6. For multivariate decoding, shaft 1 showed a *large* effect size, whereas shafts 2, 3, 4 and 6, showed a *medium* effect size. A *small* effect size was found for shaft 5. *(**B**).* Effect size (Cohen’s d) of correct minus incorrect classifications. A large effect size (Cohen’s d > .8) was found from 100 to 160 ms after stimulus onset in the alpha band (10 Hz), and from 130 to 150 ms post-stimulus onset in the high-gamma band (100-140 Hz). A medium effect size (Cohen’s d = .55) was found 200-220 ms after stimulus onset between 60 and 80 Hz, which corresponds to significant cross-correlation between decoding accuracy and correct-incorrect differences in the time-frequency domain (see Fig 4B).

***S7 Fig. Lesion data. (A).*** *MRI scan of a patient with a focal lesion of the right pSTG. (**B & C**). Psychometric functions for ba-da (**B**) and da-ta (**C**) continua. The blue line represents the linear variations of F2 (**B**), and linear variations of the voice onset time (**C**), across stimuli. Grey curves show that the patient could not discriminate between /ba/ and /da/, and identified all syllables as /da/ in the ba-da continuum, while performing normally on the da-ta continuum. Red curves indicate higher difficulty to perform the identification on the /ba/ end (**B**), but normal performance on the control da-ta task (**C**).*

***S8 Fig. Control da/ta fMRI experiment. (A).*** *Spectrograms of the stimulus continuum between syllables /da/ and /ta/, synthesized with linear variation in VOT* (-15 *ms : 5 ms :* +15 *ms). Full spectrograms at the extremes of the continuum represent /da/ and /ta/ prototype syllables (left and right panels, respectively). Middle spectrograms are centred on VOT. (**B**). Values for VOT (in blue, left panel), average d-prime (in red, middle panel), and percent syllables identified as /ta/ (in grey, right panel) (mean* ± *s.e.m.)* ***C.*** *Top panel: spatial localization of VOT neural encoding (in blue) and d-prime (in red) in fMRI bold signal, expressed as beta coefficients. Significant clusters were found in the left superior temporal gyrus (STG)* (peak MNI coordinates, *x, y, z= −66, −22,1, T = 4.2g)for the VOT-tracking, and in left inferior prefrontal (x, y, z = −48, 14, 7, T = 3.75J cortices for perceptual decision (d-prime). Images are presented at a whole-brain threshold of P<0.001. Bottom panel: percent signal change in the left inferior prefrontal cortex (in red) and in the left STG (in blue). BOLD signal increases with VOT in the left STG, and with perceptual decision load in the left inferior prefrontal region (in red).*

## References

1. Mesgarani N, Cheung C, Johnson K, Chang EF. Phonetic feature encoding in human superior temporal gyrus. Science [Internet]. 2014;343(6174):1006–10. Available from: http://www.ncbi.nlm.nih.gov/pubmed/24482117

2. Chang EF, Rieger JW, Johnson K, Berger MS, Barbaro NM, Knight RT. Categorical speech representation in human superior temporal gyrus. Nat Neurosci [Internet]. 2010 Nov [cited 2012 Oct 30];13(11):1428–32. Available from: http://www.pubmedcentral.nih.gov/articlerender.fcgi?artid=2967728&tool=pmcentrez&rendertype=abstract

3. Yan Y, Rasch MJ, Chen M, Xiang X, Huang M, Wu S, et al. Perceptual training continuously refines neuronal population codes in primary visual cortex. Nat Neurosci [Internet]. 2014;17(10):1380–7. Available from: http://dx.doi.org/10.1038/nn.3805%5Cnpapers3://publication/doi/10.1038/nn.3805

4. Hung CP, Kreiman G, Poggio T, Dicarlo JJ. Fast Readout of Object Identity from Macaque Inferior Temporal Cortex. Science. 2005;310:863–6.

5. Carota F, Kriegeskorte N, Nili H, Pulvermüller F. Representational Similarity Mapping of Distributional Semantics in Left Inferior Frontal, Middle Temporal, and Motor Cortex. 2017;1–16.

6. Fitch WT, Hauser MD. Computational Constraints on Syntactic Processing in a Nonhuman Primate. Science. 2004;303:377–80.

7. Kriegeskorte N, Kievit RA. Representational geometry: Integrating cognition, computation, and the brain. Trends Cogn Sci [Internet]. Elsevier Ltd; 2013;17(8):401–12. Available from: http://dx.doi.org/10.1016/j.tics.2013.06.007

8. Huth AG, Heer WA De, Griffiths TL, Theunissen FE, Gallant JL. Natural speech reveals the semantic maps that tile human cerebral cortex. Nature [Internet]. Nature Publishing Group; 2016;532(7600):453–8. Available from: http://dx.doi.org/10.1038/nature17637

9. Eger E, Ashburner J, Haynes J-D, Dolan RJ, Rees G. fMRI activity patterns in human LOC carry information about object exemplars within category. J Cogn Neurosci. 2008;20(2):356–70.

10. Anderson ML, Oates T. A critique of multi-voxel pattern analysis. Proc 32nd Annu Meet Cogn Sci Soc. 2010;(July):1511–16.

11. Dilks DD, Julian JB, Paunov AM, Kanwisher N. The Occipital Place Area Is Causally and Selectively Involved in Scene Perception. J Neurosci. 2013;33(4):1331–6.

12. Anderson ML. Précis of after phrenology: neural reuse and the interactive brain. Behav Brain Sci [Internet]. 2015;16:1–22. Available from: http://eprints.soton.ac.uk/252625/2/bbs.html

13. Shamma S, Lorenzi C. On the balance of envelope and temporal fine structure in the encoding of speech in the early auditory system. J Acoust Soc Am [Internet]. 2013;133(5):2818–33. Available from: http://www.pubmedcentral.nih.gov/articlerender.fcgi?artid=3663870&tool=pmcentrez&rendertype=abstract

14. Moon IJ, Won JH, Park M-H, Ives DT, Nie K, Heinz MG, et al. Optimal Combination of Neural Temporal Envelope and Fine Structure Cues to Explain Speech Identification in Background Noise. J Neurosci [Internet]. 2014;34(36):12145–54. Available from: http://www.jneurosci.org/cgi/doi/10.1523/JNEUROSCI.1025-14.2014

15. Formisano E, De Martino F, Bonte M, Goebel R. “Who” is saying “What”? Brain-based decoding of human voice and speech. Science. 2008;322:970–3.

16. Ley A, Vroomen J, Formisano E. How learning to abstract shapes neural sound representations. Front Neurosci [Internet]. 2014;8(June):1–11. Available from: http://journal.frontiersin.org/article/10.3389/fnins.2014.00132/abstract

17. Pasley BN, Knight RT. Decoding Speech for Understanding and Treating Aphasia. Prog Brain Res. 2013;207:435–56.

18. Chevillet MA, Jiang X, Rauschecker JP, Riesenhuber M. Automatic phoneme category selectivity in the dorsal auditory stream. J Neurosci. 2013;33(12):5208–15.

19. Mirman D, Chen Q, Zhang Y, Wang Z, Faseyitan OK, Coslett HB, et al. Neural organization of spoken language revealed by lesion–symptom mapping. Nat Commun [Internet]. Nature Publishing Group; 2015;6:6762. Available from: http://www.nature.com/doifinder/10.1038/ncomms7762

20. Tsunada J, Liu ASK, Gold JI, Cohen YE. Causal contribution of primate auditory cortex to auditory perceptual decision-making. Nat Neurosci [Internet]. 2016;19(1):135–42. Available from: http://www.nature.com/neuro/journal/v19/n1/full/nn.4195.html?WT.ec_id=NEURO-201601&spMailingID=50354006&spUserID=Njk2Njk2MzE4MjUS1&spJobID=824237578&spReportId=ODI0MjM3NTc4S0

21. Zatorre RJ, Belin P. Spectral and temporal processing in human auditory cortex. Cereb Cortex [Internet]. 2001;11(10):946–53. Available from: http://www.ncbi.nlm.nih.gov/pubmed/11549617

22. Fontolan L, Morillon B, Liegeois-Chauvel C, Giraud A-L. The contribution of frequency-specific activity to hierarchical information processing in the human auditory cortex. Nat Commun [Internet]. Nature Publishing Group; 2014;5(May):4694. Available from: http://www.nature.com/doifinder/10.1038/ncomms5694

23. Arnal LH, Giraud A-L. Cortical oscillations and sensory predictions. Trends Cogn Sci [Internet]. Elsevier Ltd; 2012 Jul [cited 2012 Oct 30];16(7):390–8. Available from: http://www.ncbi.nlm.nih.gov/pubmed/22682813

24. Fries P. Rhythms for Cognition: Communication through Coherence. Neuron [Internet]. Elsevier Inc.; 2015;88(1):220–35. Available from: http://linkinghub.elsevier.com/retrieve/pii/S0896627315008235

25. Donoso M, Collins AGE, Koechlin E. Human cognition. Foundations of human reasoning in the prefrontal cortex. Science [Internet]. 2014;344(6191):1481–6. Available from: http://www.ncbi.nlm.nih.gov/pubmed/24876345

26. Blumstein SE, Myers EB, Rissman J. The perception of voice onset time: an fMRI investigation of phonetic category structure. J Cogn Neurosci [Internet]. 2005 Sep;17(9):1353–66. Available from: http://www.ncbi.nlm.nih.gov/pubmed/16197689

27. Rogers JC, Davis MH. Inferior Frontal Cortex Contributions to the Recognition of Spoken Words and Their Constituent Speech Sounds. J Cogn Neurosci XY,. 2017;27:1–18.

28. Marti S, King J-R, Dehaene S. Time-Resolved Decoding of Two Processing Chains during Dual-Task Interference. Neuron [Internet]. 2015;88:1297–307. Available from: http://linkinghub.elsevier.com/retrieve/pii/S0896627315009344

29. King J-R, Dehaene S. Characterizing the dynamics of mental representations: the temporal generalization method. Trends Cogn Sci [Internet]. Elsevier Ltd; 2014;18(4):203–10. Available from: http://linkinghub.elsevier.com/retrieve/pii/S1364661314000199

30. Isik L, Meyers EM, Leibo JZ, Poggio T. The dynamics of invariant object recognition in the human visual system. J Neurophysiol [Internet]. 2014;111(1):91–102. Available from: http://www.ncbi.nlm.nih.gov/pubmed/24089402

31. Du Y, Buchsbaum BR, Grady CL, Alain C. Noise differentially impacts phoneme representations in the auditory and speech motor systems. Proc Natl Acad Sci U S A [Internet]. 2014;111(19):7126–31. Available from: http://www.ncbi.nlm.nih.gov/pubmed/24778251

32. Liebenthal E, Sabri M, Beardsley S a, Mangalathu-Arumana J, Desai A. Neural dynamics of phonological processing in the dorsal auditory stream. J Neurosci [Internet]. 2013;33(39):15414–24. Available from: http://www.pubmedcentral.nih.gov/articlerender.fcgi?artid=3782621&tool=pmcentrez&rendertype=abstract

33. Obleser J, Eisner F, Kotz S a. Bilateral speech comprehension reflects differential sensitivity to spectral and temporal features. J Neurosci [Internet]. 2008 Aug 6 [cited 2012 Nov 12];28(32):8116–23. Available from: http://www.ncbi.nlm.nih.gov/pubmed/18685036

34. Liegeois-Chauvel C, Giraud K, Badier J-M, Marquis P, Chauvel P. Intracerebral evoked potentials in pitch perception reveal a function of asymmetry of human auditory cortex. Ann N Y Acad Sci. 2001;930:117–32.

35. Alain C, Snyder JS. Age-related differences in auditory evoked responses during rapid perceptual learning. Clin Neurophysiol. 2008;119(2):356–66.

36. Alain C, Snyder JS, He Y, Reinke KS. Changes in auditory cortex parallel rapid perceptual learning. Cereb Cortex. 2007;17(5):1074–84.

37. Binder JR, Liebenthal E, Possing ET, Medler D a, Ward BD. Neural correlates of sensory and decision processes in auditory object identification. Nat Neurosci. 2004;7(3):295–301.

38. Giraud A-L, Kell C, Thierfelder C, Sterzer P, Russ M, Preibisch C, et al. Contributions of sensory input, auditory search and verbal comprehension to cortical activity during speech processing. Cereb Cortex. 2004;14(3):246–55.

39. Lee Y-S, Turkeltaub P, Granger R, Raizada RDS. Categorical speech processing in Broca’s area: an fMRI study using multivariate pattern-based analysis. J Neurosci [Internet]. 2012 Mar 14 [cited 2012 Nov 10];32(11):3942–8. Available from: http://www.ncbi.nlm.nih.gov/pubmed/22423114

40. Arsenault JS, Buchsbaum BR. Distributed Neural Representations of Phonological Features during Speech Perception. J Neurosci. 2015;35(2):634–42.

41. Blank H, Davis MH. Prediction Errors but Not Sharpened Signals Simulate Multivoxel fMRI Patterns during Speech Perception. PLoS Biol [Internet]. 2016;14(11):e1002577. Available from: http://www.ncbi.nlm.nih.gov/pubmed/27846209

42. Bonte M, Hausfeld L, Scharke W, Valente G, Formisano E. Task-Dependent Decoding of Speaker and Vowel Identity from Auditory Cortical Response Patterns. J Neurosci. 2014;34(13):4548 –4557.

43. Sohoglu E, Peelle JE, Carlyon RP, Davis MH. Predictive top-down integration of prior knowledge during speech perception. J Neurosci [Internet]. 2012 Jun 20 [cited 2012 Oct 26];32(25):8443–53. Available from: http://www.ncbi.nlm.nih.gov/pubmed/22723684

44. Kok P, Jehee JFM, de Lange JFP. Less Is More: Expectation Sharpens Representations in the Primary Visual Cortex. Neuron [Internet]. 2012;75(2):265–70. Available from: http://linkinghub.elsevier.com/retrieve/pii/S0896627312004382

45. Weichwald S, Meyer T, Özdenizci O, Schölkopf B, Ball T, Grosse-Wentrup M. Causal interpretation rules for encoding and decoding models in neuroimaging. Neuroimage [Internet]. Elsevier Inc.; 2015;110:48–59. Available from: http://dx.doi.org/10.1016/j.neuroimage.2015.01.036

46. Zohary E, Celebrini S, Britten KH, Newsome WT. Neuronal plasticity that underlies improvement in perceptual performance. Science. 1994;263:1289–92.

47. Sompolinsky H, Yoon H, Kang K, Shamir M. Population coding in neuronal systems with correlated noise. Phys Rev E Stat Nonlin Soft Matter Phys. 2001;64:51904.

48. Pitkow X, Liu S, Angelaki DE, DeAngelis GC, Pouget A. How Can Single Sensory Neurons Predict Behavior? Neuron [Internet]. Elsevier Inc.; 2015;87(2):411–23. Available from: http://linkinghub.elsevier.com/retrieve/pii/S0896627315005966

49. Çukur T, Nishimoto S, Huth AG, Gallant JL. Attention during natural vision warps semantic representation across the human brain. Nat Neurosci [Internet]. Nature Publishing Group; 2013;16(6):763–70. Available from: http://www.nature.com.ezproxy.cul.columbia.edu/neuro/journal/v16/n6/full/nn.3381.html

50. Cheung C, Hamiton LS, Johnson K, Chang EF. The auditory representation of speech sounds in human motor cortex. Elife [Internet]. 2016;5:1–19. Available from: http://elifesciences.org/lookup/doi/10.7554/eLife.12577

51. Correia J, Formisano E, Valente G, Hausfeld L, Jansma B, Bonte M. Brain-Based Translation: fMRI Decoding of Spoken Words in Bilinguals Reveals Language-Independent Semantic Representations in Anterior Temporal Lobe. J Neurosci [Internet]. 2014;34(1):332–8. Available from: http://www.jneurosci.org/cgi/doi/10.1523/JNEUROSCI.1302-13.2014

52. Kawahara H, Nisimura R, Irino T, Morise M, Takahashi T, Banno H. Temporally variable multi-aspect auditory morphing enabling extrapolation without objective and perceptual breakdown. In: ICASSP, IEEE International Conference on Acoustics, Speech and Signal Processing - Proceedings. 2009. p. 3905–8.

53. Kroon P, Kleijn WB. Linear-prediction based analysis-by-synthesis coding. Elsevier S. 1995. 79–119 p.

54. Riede T, Suthers RA. Vocal tract motor patterns and resonance during constant frequency song: The white-throated sparrow. J Comp Physiol A Neuroethol Sensory, Neural, Behav Physiol. 2009;195:183–92.

55. Goncharoff V, Kaine-Krolak M. Interpolation of LPC spectra via pole shifting. Ieee Int Conf Acoust Speech Signal Process [Internet]. 1995;1:780–780.

56. Tadel F, Baillet S, Mosher JC, Pantazis D, Leahy RM. Brainstorm: A user-friendly application for MEG/EEG analysis. Comput Intell Neurosci. Hindawi Publishing Corporation; 2011;2011.

57. Oostenveld R, Fries P, Maris E, Schoffelen JM. FieldTrip: Open source software for advanced analysis of MEG, EEG, and invasive electrophysiological data. Comput Intell Neurosci. Hindawi Publishing Corporation; 2011;2011.

58. Liebenthal E, Binder JR, Spitzer SM, Possing ET, Medler D a. Neural substrates of phonemic perception. Cereb Cortex [Internet]. 2005 Oct [cited 2012 Nov 18];15(10):1621–31. Available from: http://www.ncbi.nlm.nih.gov/pubmed/15703256

59. Hämäläinen M, Ilmoniemi R. Interpreting magnetic fields of the brain: minimum norm estimates. Med Biol Eng Comput. 1994;32(1):35–42.

60. Destrieux C, Fischl B, Dale A, Halgren E. Automatic parcellation of human cortical gyri and sulci using standard anatomical nomenclature. Neuroimage. 2010;53:1–15.

61. Dhamala M, Rangarajan G, Ding M. Analyzing information flow in brain networks with nonparametric Granger causality. Neuroimage. 2008;41(2):354–62.

62. Dhamala M, Rangarajan G, Ding M. Estimating granger causality from fourier and wavelet transforms of time series data. Phys Rev Lett. 2008;100(1):1–4.

63. Lyu B, Ge J, Niu Z, Tan LH, Gao J-H. Predictive Brain Mechanisms in Sound-to-Meaning Mapping during Speech Processing. J Neurosci. 2016;36(42):10813–22.

64. Yoo S, Lee K-M. Articulation-based sound perception in verbal repetition: a functional NIRS study. Front Hum Neurosci [Internet]. 2013;7(September):540. Available from: http://www.pubmedcentral.nih.gov/articlerender.fcgi?artid=3763229&tool=pmcentrez&rendertype=abstract

65. Klein M, Grainger J, Wheat KL, Millman RE, Simpson MIG, Hansen PC, et al. Early Activity in Broca’s Area during Reading Reflects Fast Access to Articulatory Codes from Print. Cereb Cortex. 2015;25(7):1715–23.

66. Meyers EM. The Neural Decoding Toolbox. Front Neuroinform [Internet]. 2013;7:8. Available from: http://www.pubmedcentral.nih.gov/articlerender.fcgi?artid=3660664&tool=pmcentrez&rendertype=abstract

67. Maris E, Oostenveld R. Nonparametric statistical testing of EEG- and MEG-data. J Neurosci Methods [Internet]. 2007 Aug 15 [cited 2012 Oct 29];164(1):177–90. Available from: http://www.ncbi.nlm.nih.gov/pubmed/

